# Effects of seasonality and land use on the abundance and distribution of mosquitoes on St. Kitts, West Indies

**DOI:** 10.1101/2020.05.11.089037

**Authors:** Matthew J. Valentine, Brenda Ciraola, Gregory R. Jacobs, Charlie Arnot, Patrick J. Kelly, Courtney C. Murdock

## Abstract

**Background:** High quality mosquito surveys that collect fine resolution local data on mosquito species’ abundances provide baseline data to help us understand potential host-pathogen-mosquito relationships, accurately predict disease transmission, and target mosquito control efforts in areas at risk of mosquito borne diseases.

**Methods:** As part of an investigation into arboviral sylvatic cycles on the Caribbean island of St. Kitts, we carried out an island wide mosquito survey from November 2017 to March 2019. Using Biogents Sentinel 2 and miniature CDC light traps that were set monthly and run for 48 hour intervals, we collected mosquitoes from a total of 30 sites distributed across the five common land covers on the island (agricultural, mangrove, rainforest, scrub, and urban). We developed a mixed effects negative binomial regression model to predict the effects of land cover, seasonality, and precipitation on observed counts of the most abundant mosquito species we found.

**Results:** We captured 10 of the 14 mosquito species reported on the island, the four most abundant being *Aedes taeniorhynchus, Culex quinquefasciatus, Aedes aegpyti*, and *Deinocerites magnus*. Sampling in the mangroves yielded the most mosquitoes, with *Ae. taeniorhynchus, Cx. quinquefasciatus*, and *De. magnus* predominating. *Aedes aegypti* was recovered primarily from urban and agricultural habitats, but also at lower frequency in other land covers. *Psorophora pygmaea* and *Toxorhynchites guadeloupensis* were only captured in scrub habitat. Capture rates in rainforests were low. Our models indicated the relative abundance of the four most common species varied seasonally and with land cover. They also suggested that the extent to which monthly average precipitation influenced counts varied according to species.

**Conclusions:** This study demonstrates there is high seasonality in mosquito abundances and that land cover influenced the distribution and abundance of mosquito species on St. Kitts. Further, human-adapted mosquito species (e.g. *Ae. aegypti* and *Cx. quinquefasciatus*) that are known vectors for many human relevant pathogens are the most wide-spread (across land covers) and the least responsive to seasonal variation in precipitation.

## Background

Mosquitoes are responsible for considerable human and animal suffering and economic losses because of their nuisance value and the diseases of high morbidity and mortality they can transmit [1–2]. Recent mosquito-borne arboviral pandemics have been able to spread through previously unaffected regions, like the Americas [3–4], due to the wide-spread presence and abundance of human-adapted mosquito species. Furthermore, mosquito-borne pathogens can become established in new areas if there are suitable animal reservoir populations and mosquito species that can transmit the organisms between these animals (and potentially to humans), as has occurred with yellow fever virus in South America [5–6].

High quality mosquito surveys are an essential tool for predicting mosquito-borne disease transmission and for mosquito control. Surveys that collect fine resolution local data on mosquito species abundances provide fundamental baseline data on the composition of mosquito communities in a given area, the relative abundances of mosquito species within the community, and how the abundance of species and the composition of mosquito communities change across space and time. The development of population abundance models that leverage count data generated from these surveys, in turn, can be used to predict how mosquito abundances change seasonally and across different land covers. Information of this nature is crucial for describing potential host-pathogen-mosquito relationships in novel transmission foci, accurately predicting disease transmission, and for targeting and assessing the efficacy of mosquito control efforts [7–8].

St. Kitts is a small tropical island in the Caribbean where local experience shows mosquitoes are very common and their nuisance value high. Outbreaks of mosquito-borne arboviral infections have occurred on the island, which also has a large population of African green monkeys (*Chlorocebus aethiops sabaeus*) that may be involved in arbovirus sylvatic cycles as is the case in Africa [6]. Due to the large numbers of tourists visiting the region each year, islands in the Caribbean like St. Kitts could be a source of mosquitoes, and the pathogens they carry, for transfer into currently naïve areas of the world like the USA. Indeed, some workers consider islands, ‘hotspots’ for disease emergence [9].

Historically, there has been long standing interest in the mosquito species inhabiting the Caribbean particularly since it was discovered that malarial parasites (*Plasmodium* spp.) and yellow fever virus are transmitted by mosquitoes. Detailed mosquito surveys from the 1970s included several Caribbean islands including St. Kitts and Nevis [10]. The most recent survey on St. Kitts was conducted in 2010 [11] and although this was the most comprehensive survey performed on the island to date, it did not provide data on the how the distribution and relative abundances of mosquito species changes seasonally and with land cover. Mosquitoes were only collected during a single week of the dry and wet seasons and sampling did not include all land covers. As part of an investigation into arboviral sylvatic cycles on St. Kitts, we carried out a comprehensive survey of the mosquito populations across the various land covers on the island on a monthly/ bimonthly basis from September 2017 to March 2019. We related mosquito survey data to relevant biological and environmental covariates to assess the influence of land use and seasonal variation on spatial and temporal biodiversity of mosquitoes on St. Kitts. Below are a description of our methods and our findings.

## Methods

### Study Area

St. Kitts (Fig. 1) is a 168 square km, geographically isolated, volcanic, Caribbean island located in the Lesser Antilles (17.33° N, 62.75° W). It has a population of approximately 40,000 people mostly inhabiting Basseterre, the capital, and a string of small village communities distributed along the main coastal road which circles the island. The climate in St. Kitts is tropical, driven by constant sea breezes with little seasonal temperature variation (27-30°C). The wet season runs from May to November with risk of hurricanes from June to November. Rainforest covers the uninhabited, steep volcanic slopes in the center of the island, surrounded by lower gentler slopes consisting mostly of abandoned sugar cane fields or arable farmlands. The south east of the island is primarily an arid peninsula covered mainly in scrub with beaches, mangroves, and salt-ponds.

**Fig 1.**
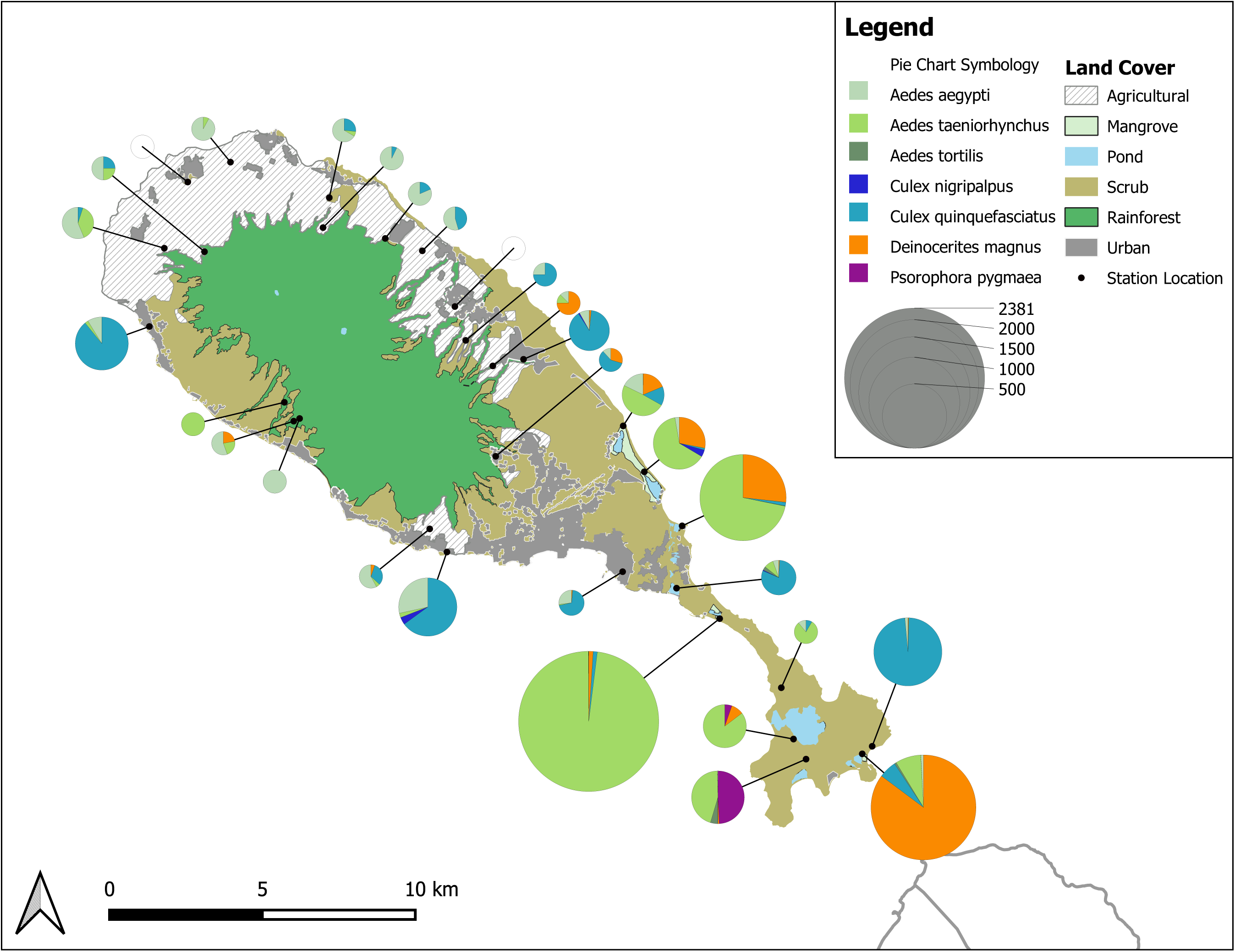
Numbers and species of mosquitoes trapped on St. Kitts at 30 sites comprising six replicates in each of five land covers

### Mosquito Sampling

To estimate the diversity and abundance of mosquito species across different land covers and seasons, we evaluated counts and species identity of mosquitoes captured in trap arrays set across the island. Trapping was carried out once every one to two months from September 2017 to March 2019 in each of five representative land covers (September and October 2017 were omitted from the analysis due to severe weather). We used a randomized stratified sampling design to select six replicate sites at least 1 km apart, in each of five distinct land cover categories (agricultural, mangrove, rainforest, scrub and urban) producing 30 sites in total (Fig. 1). Final site selection was dependent on accessibility and landowner consent.

To increase detection of host-seeking mosquitoes we used both CDC light traps (J.W. Hock, USA) and Biogents Sentinel 2 traps (BGS) (Biogents AG, Germany) until November 2018, thereafter only BGS traps were used. The CDC light trap was run without the light source to decrease potential non-mosquito bycatch, and the BG-sentinel lure (Biogents AG, Germany) was used in the BGS trap to target anthrophilic species. Carbon dioxide was generated by mixing 35 g of dried bread making yeast (Fleischmann’s Active Dry Yeast, USA), 0.7 kg of unbranded white sugar, and approximately 2.5 L water in a centrally placed 5 L water bottle. Carbon dioxide was delivered to each trap via a 5 m length of 5 mm (internal diameter) PVC tubing [12–14]. At each site, traps were placed 10 m apart from one another. Traps were run each month when possible from September 2017 to March 2019 for 48 hours, with yeast-sugar solution, batteries, and catch bags replaced every 24 hours. Trapped mosquitoes were transported to the research laboratory of Ross University School of Veterinary Medicine (RUSVM) and stored at - 80°C for later identification. After being rehydrated on chilled damp tissue paper, mosquitoes were identified on a chill table using morphological keys under a stereomicroscope (Cole Palmer, USA) at 10x-40x magnification [15–17]. Counts of each mosquito species were recorded for each sampling date, land use, and location.

### Statistical Analysis

We evaluated influences of different land covers (agricultural, mangrove, rainforest, scrub, urban) on the relative abundance of the four most common mosquito species found in our survey. The land covers in a 1 km^2^ area (565 m radius) around each sampling site were determined from local observation, a remote sensing vegetation classification [18], assorted vector data sets supplied by the government of St. Kitts, and the most recent Google Imagery (2019). When discordance in ascribing land covers was found between the different methods, the Google images were used preferentially. The percentages of each land cover at each site (Fig. 2) were calculated and used in establishing the models.

**Fig 2.**
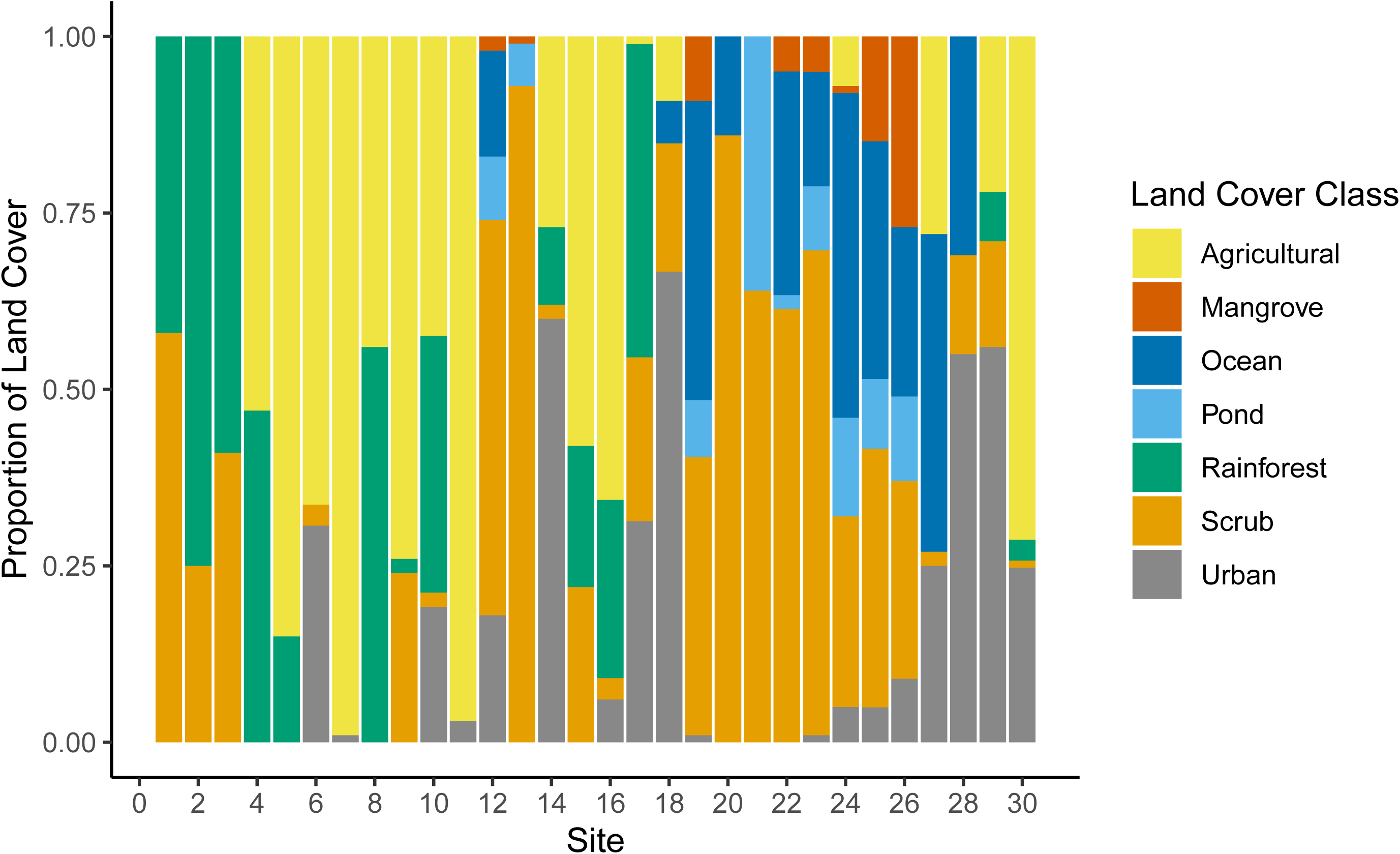
The proportions of the different land covers found in a 1 km^2^ area (565 m radius) around each of the trapping 30 sites used in the study

We assessed the effects of land cover at each trap location and monthly precipitation on the numbers of the four most common mosquito species trapped using mixed effects generalized linear regression models [19]. Our response variable for this analysis was the number of mosquitoes of each species captured by BGS traps during each 48 hour trapping interval. We omitted CDC traps in the model because they were not used on all occasions and seemed to capture fewer mosquitoes. A list of variables and general expectations of their effects on the counts of mosquitoes of each species captured can be found in Additional file 1: Table S1. We included two categorical variables reflecting the high affinity of some mosquito species (e.g. *Ae. taeniorhynchus* and *De. magnus*) for mangrove habitat and crabhole habitat located in the vicinity of mangroves [10, 20–22]. These variables included “mangrove”, which described the land cover of sites that fell within mangrove habitats regardless of surrounding land covers, and “m_trait”, which described a mosquito species preference for mangrove habitat. Monthly precipitation measurements were obtained at the Robert L. Bradshaw International Airport and accessed as archived data downloaded from the Weather Underground website (www.wunderground.com: accessed August 2019).

We fitted models using the R package glmmTMB, which allows the specification of generalized linear mixed-effects models for a variety of error distributions, including Poisson and negative binomial distributions [23]. Preliminary analyses revealed that a negative binomial distribution with a quadratic variance-to-mean relationship best explained our data [24] (Additional file 2: Table S2), and we used this error distribution for all subsequent analyses. We assumed a linear relationship between overall mosquito counts (on the log-link scale) and monthly average precipitation at the island scale to account for intra-annual seasonality (e.g., wet vs. dry seasons). Species-specific random slope and intercept terms for precipitation allow its effect to vary by species, and random site intercepts account for repeat observations at each site. The effects of land cover were allowed to vary independently by species. We evaluated 12 model hypotheses of species-specific variation in relative abundance with land cover (Table 1). Land cover variables incorporated in our 12 models generally include the percentage of local land cover. We excluded any hypotheses for which variance inflation factors were greater than five prior to model evaluation and used AIC to select our best model from the candidate set [25]. To assess model fit, we evaluated the distribution of re-scaled model residuals from the R package DHARMa [26] and calculated conditional and marginal *R*^*2*^ values following Nakagawa et al [27]. We used site-level predictions from our best model to show model-estimated trends in mosquito abundance across land use and season for the duration of our study.

**Table 1.** A list of main effects of all candidate models considered in analyses of land cover effects on mosquito relative abundance. Interaction-only terms only were used between indicator variables, and evaluates the effect when both indicator variables are equal to 1 (i.e., “m_trait: Mangrove” evaluates the effect of having the mangrove breeding trait when located in a mangrove)

| Name | Main Effects | ΔAIC | K |
| --- | --- | --- | --- |
| H1 | Precip + LocalAgriculture + LocalUrban + LocalRainforest + LocalMangrove + spp<sup>a</sup>Mangrove | 11.4 | 26 |
| H2 | Precip + LocalAnthropogenic | 53.6 | 11 |
| H3 | Precip + LocalAgriculture + LocalUrban | 31 | 15 |
| H4 | Precip + LocalAnthropogenic + m_trait'Mangrove | 38.6 | 14 |
| H5 | Precip + LocalAgriculture + LocalUrban + m_trait'Mangrove | 14.5 | 18 |
| H6 | Precip + LocalScrub+ LocalAnthropogenic + m_trait'Mangrove | 37.2 | 18 |
| H7 | Precip + LocalAgriculture + LocalUrban + LocalScrub+ m_trait'Mangrove | 15 | 22 |
| H8 | Precip + LocalRainforest + LocalAnthropogenic + m_trait'Mangrove | 25.6 | 18 |
| H9 | Precip + LocalAgriculture + LocalUrban + LocalRainforest + m_trait'Mangrove | 3.9 | 20 |
| H10 | Precip + LocalRainforest + LocalScrub+ LocalAnthropogenic + m_trait'Mangrove | 26.8 | 22 |
| H11 | Precip + LocalAgriculture + LocalUrban + LocalRainforest + LocalScrub+ m_trait'Mangrove | 3.9 | 26 |
| H12 | Precip + LocalAgriculture + LocalUrban + LocalRainforest + m_trait<sup>a</sup>Mangrove | 0 | 22 |

<sup>a</sup>Two variables with full factorial interaction comprised of main effects and interaction terms : cases where only the interaction term is evaluated

## Results

From November 2017 to March 2019 we captured 10 of the 14 species previously recorded on St. Kitts [10–11, 15] (Additional file 3: Table S3, Additional file 4: Table S4). Voucher specimens were deposited in the United States National Museum (USNM) under the following catalog numbers USNMENT01239050-74 and USNMENT01239079. The most abundant species of mosquito was *Aedes taeniorhynchus* (n=3861), which was primarily found in mangroves (88.4%) (Figure 1, Additional file 3: Table S3). *Culex quinquefasciatus* (n=1663) was the second most abundant species primarily captured in urban areas (48.8%) (Figure 1, Additional file 3: Table S3). *Aedes aegypti, Ae. taeniorhynchus, Cx. quinquefasciatus*, and *Deinocerites magnus* were species captured in all five land covers (Figure 1, Additional file 3: Table S3). There were no clear morphological differences in the *Ae. aegypti* collected across different land covers. All other species were much less abundant. *Psorophora pygmaea* and *Toxorhynchites guadeloupensis* were only captured in scrub habitat, with the remaining species being distributed across more than one land cover (Figure 1, Additional file 3: Table S3). The highest overall number of mosquitoes captured, mostly *Ae. taeniorhynchus*, were caught in November 2018 (N=3786), and monthly mean average catches were higher in general during the wet season (N=1080) than the dry season (N=177). *Aedes aegypti* was the only species to be captured during every trapping month. Only four *Anopheles albimanus*, the main vector of malaria in the Caribbean [28], were caught across the entire survey period (Additional file 4: Table S4).

The best model of temporal variation in the relative abundance of the four most common mosquito species on the Island of St. Kitts was the model H12 from Table 1. This model included the effects of monthly precipitation, a mangrove breeding trait interaction with mangrove sites, and species-specific effects driven by agriculture, urban, and rainforest land covers (Table 2). Our best model can be expressed in pseudo-code as

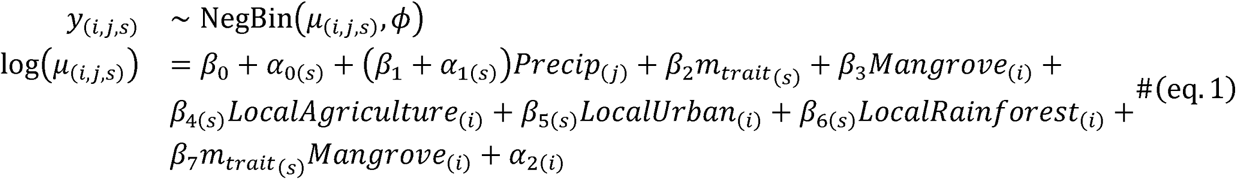

where *i, j*, and denote indices for site, month, and species, respectively. NegBin reflects the negative binomial distribution, and the final model includes the following: random intercepts for each species (*β*_0_ + *α*_0(*s*)_), a main effect of precipitation with species-specific random slopes ((*β* _l_ + *α* _l(*s*)_)*Precip*_(*j*)_), an interaction between mangrove and the mangrove breeding trait (*β* _2_*m*_*trait*_(*s*)_ + *β* _3_*Mangrove*_(*i*)_ + *β* _7_*m*_*trait* _(*s*)_*Mangrove*_(*i*)_), species-specific main effects on proportional local land cover variables (*β* _4(*s*)_ *LogalAgriculture e*_(*i*)_ + *β*_5(*s*)_ *LocalUrban* _(*i*)_ + *β* _6(*s*)_ *LocalRainforest*_(*i*)_), and a site-level random intercept term (*α* _2(*i*)_). The variance for our best negative binomial model scales quadratically with the mean (µ): 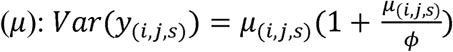 [24]. Our best model’s predicted relative abundance for each mosquito species, averaged across each site’s land cover, illustrated species-specific responses to surrounding land cover and seasonal precipitation effects across the island (Table 2, Figure 3, Additional file 5: Figure S1). Overall, we observed a positive effect of precipitation on the relative abundance of mosquitoes (*β* _l_), with significant among-species variation in this relationship (*α* _l(*s*)_, Table 2). To further explore the effect of precipitation, we derived the best linear unbiased predictors (BLUPs) for each species’ precipitation effect. The BLUPs suggested that the effect of precipitation was strongest for *Ae. taeniorhynchus* and to a lesser extent *Ae. aegypti* (Figure 4), and was weakest for *Cx. quinquefasciatus* and *De. magnus*. The model also predicted significantly negative relationships between urban land cover and *Ae. taeniorhynchus* and *De. magnus*, but only slightly-positive, non-significant relationships between urban land cover and the urban-associated mosquitoes, *Cx. quinquefasciatus* and *Ae. aegypti*. Rainforest and agricultural land cover were negatively associated with the relative abundance of all species we considered except *Ae. aegypti*, which exhibited no significant covariation with either rainforest or agricultural land cover (Table 2). As expected, the model predicts the relative abundance of mosquito species with a mangrove breeding preference (*Ae. taeniorhynchus* and *De. magnus*) to be lower than average except when trapping sites occur within mangrove habitat where the mangrove “trait” had a net positive effect (i.e., (*β* _2_ + *β* _7_)> 0, Table 2). Inspection of re-scaled residuals generated by simulation from the fitted model [26] suggested uniformity in the distribution of residuals (One-sample Kolmogorov-Smirnov test: *D* = 0.018, *P* = 0.671), indicating good concurrence between the data and model predictions. Marginal and conditional *R*^2^ (0.601 and 0.696, respectively) also indicated a well-fit model (Table 2): the marginal *R*^2^ of 0.601 suggested that the fixed effects portion of the model explained over 60% of the variation in counts, and the conditional *R*^2^ of 0.695 revealed the additional variance explained by accounting for additional variation attributable to the random effects [27].

**Table 2.**
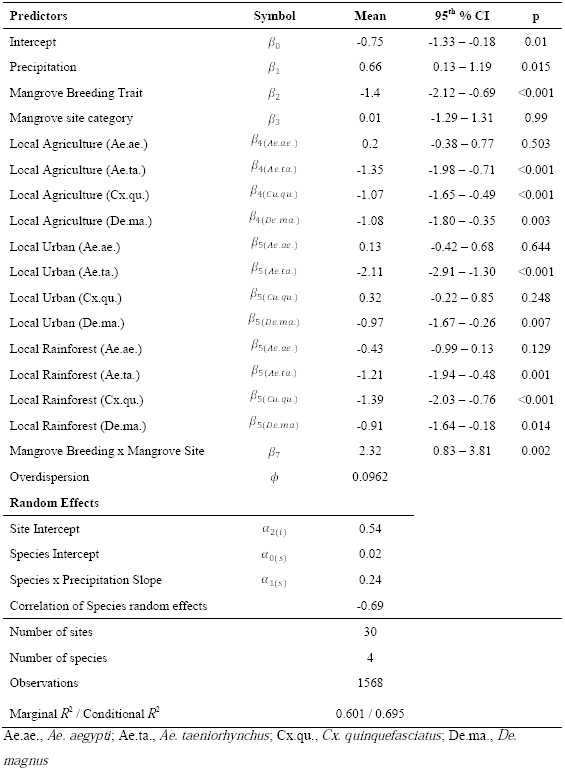
Model parameters and diagnostics from our best model of mosquito counts. Means and confidence intervals are given on the log-scale, and symbols correspond to parameters in equation 1

**Fig 3.**
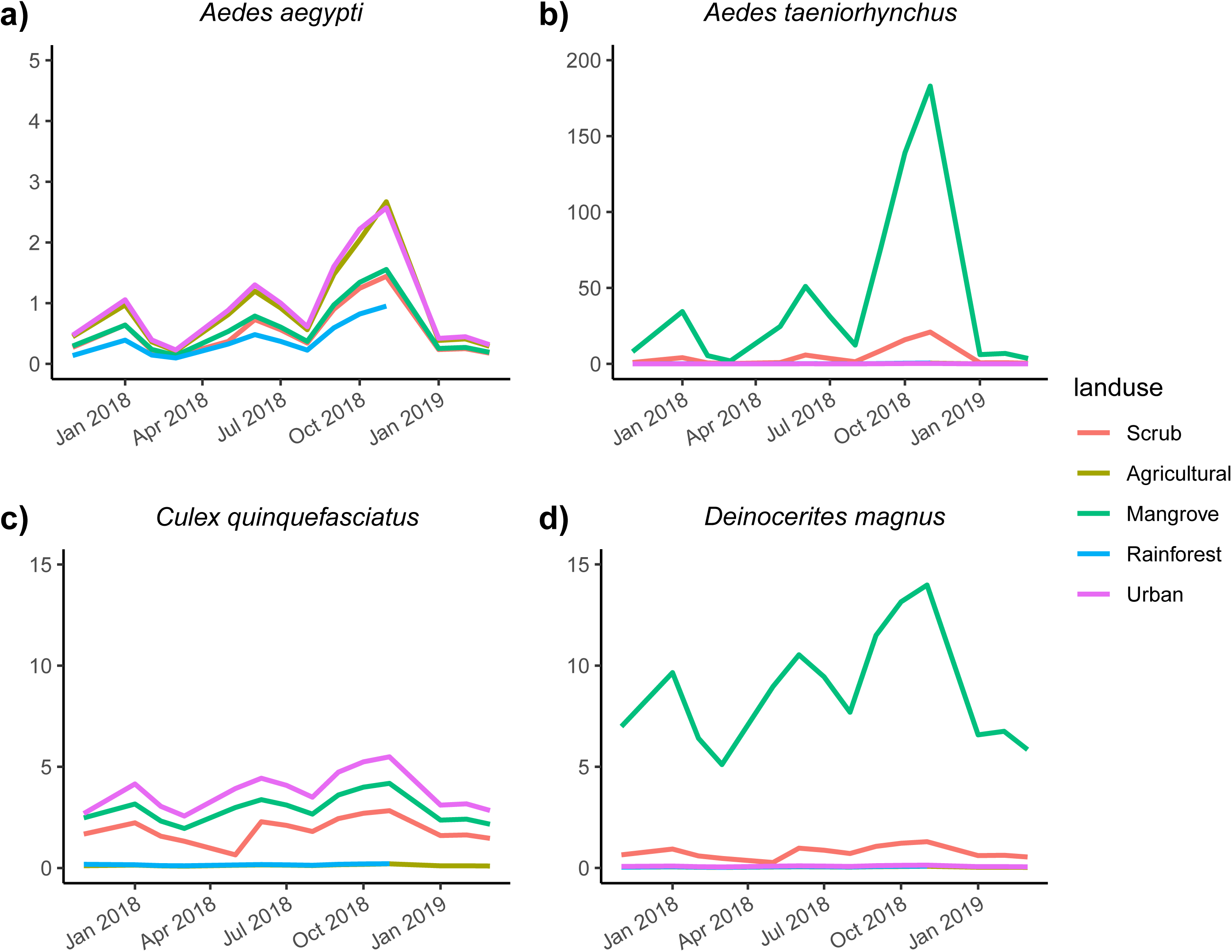
Time series plot for each site land cover (line color) and for the four major species we captured (panel) in their predicted relative abundance (conditional on random effects) from our best model. Solid lines denote the average predicted relative abundance of mosquito species across the 6 sites located within that land cover

**Fig 4.**
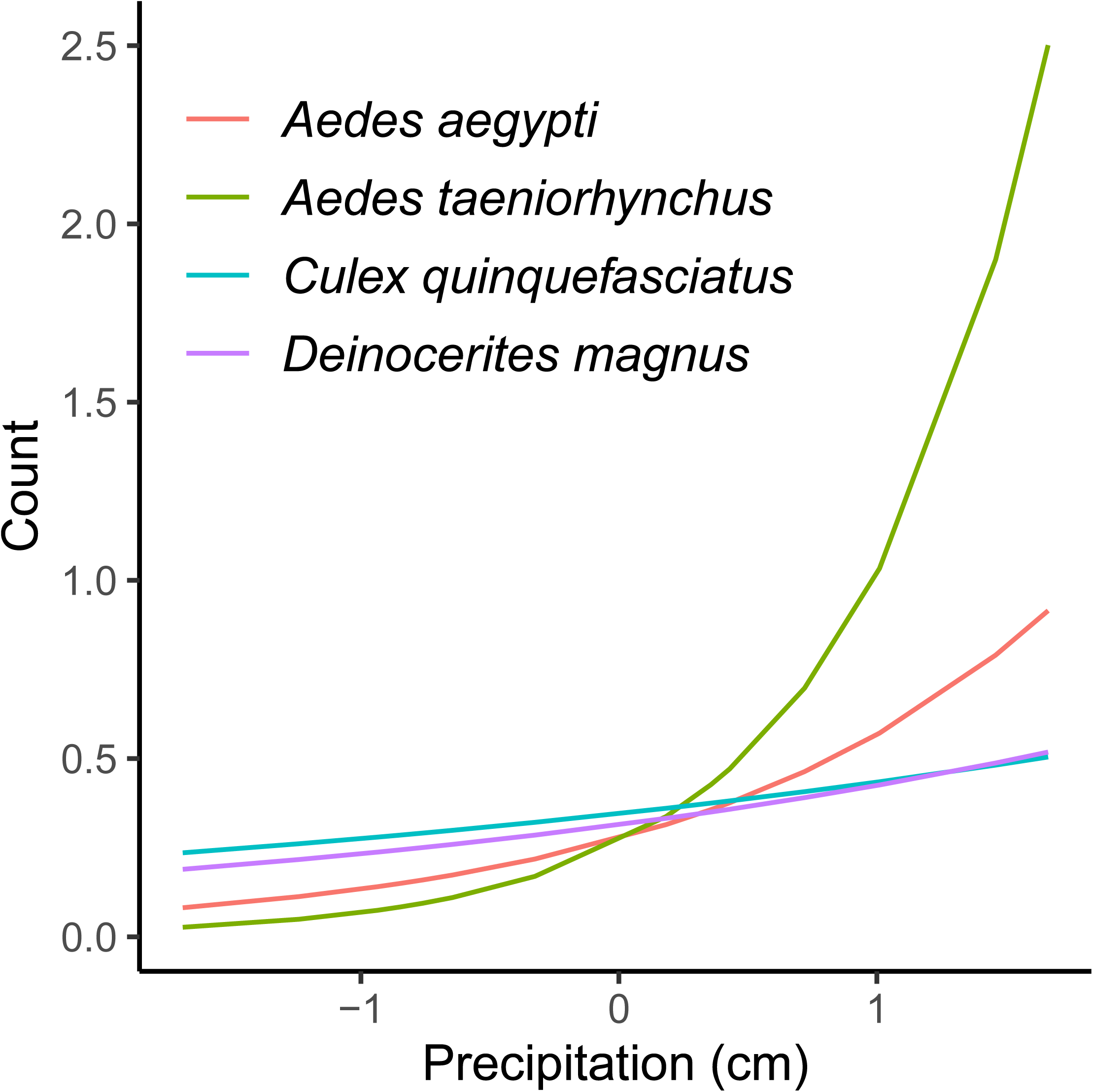
Best linear unbiased predictors (BLUPs) for the effect of precipitation on mosquito counts

## DISCUSSION

Monthly mosquito surveillance on St. Kitts from November 2017 to March 2019 enabled us to capture a diversity of mosquito species that varied in abundance across seasons and land cover types. We captured 10 of the 14 species (5 genera) historically recorded on the island [10–11, 15]. While our results largely confirm those of the 2010 survey, they provide higher spatial and temporal resolution of both the relative abundance and distribution of different mosquito species on the island.

The four most abundant species captured were *Ae. taeniorhynchus, Cx. quinquefasciatus, Ae. aegypti*, and *De. magnus*. These species were detected at least once in all land cover types over the study period. The three most abundant species captured are competent vectors of pathogens recorded on St. Kitts (*Cx. quinquefasciatus* - West Nile virus and *Dirofilaria immitis*; *Ae. taeniorhynchus* – *Dirofilaria immitis*; and *Ae. aegypti* - dengue, chikungunya, and Zika viruses) [29–33] and other pathogens that could potentially become established on the island in the future due to the abundance of their vectors. For example, *Cx. quinquefasciatus*, the southern house mosquito, is widely distributed across the sub tropics and can also transmit St. Louis encephalitis virus, Western equine encephalitis virus, Rift Valley fever virus, *Wuchereria bancrofti*, and avian malaria [17]. *Aedes taeniorhynchus*, the black salt marsh mosquito, is widely distributed in all islands of the Caribbean and can transmit Venezuelan, Eastern and Western equine encephalitis virus [11, 34–35]. The eponymous yellow fever mosquito, *Ae. aegypti*, can also transmit yellow fever virus [17]. While *An. albimanus* is a known malaria vector in Central America, northern South America, and the Caribbean, the overall low abundance of this species on St. Kitts (n=4 in our study and n=2 in 2010, [11]) suggests this species is unlikely to support malaria transmission on the island.

The presence and absence, as well as overall relative abundance, of particular mosquito species captured across the different land covers on St. Kitts broadly align with what is known for these species in the literature. The species count model for our four most abundant mosquitoes predicts both *Cx. quinquefasciatus* and *Ae. aegypti* to have the highest relative abundance in an urban habitat. Both species breed most successfully in fresh water-filled man-made containers and are therefore found primarily around houses in urban environments. Further, *Ae. aegypti* preferentially feeds on human hosts, particularly when indoors [17, 36] and rests inside domestic dwellings [17]. The fact that we observed these species, albeit at lower abundances, in other land covers is not entirely surprising. The model predicted *Ae. aegypti* to be similarly abundant across the survey period in urban as well as agricultural habitats. From local experience, agricultural land covers provide ample breeding habitats for container breeding mosquito species in the form of discarded tires, styrofoam containers, plastic water bottles and bags, and agricultural equipment in which water can collect.

Interestingly, the model also predicted moderate relative abundance across the survey period for both *Cx. quinquefasciatus* and *Ae. aegypti* in mangrove and scrub habitats, which may be due to the presence of artificial containers suitable for breeding or an increased tolerance to brackish water in high marsh areas of the mangrove [21, 37]. *Culex quinquefasciatus* has been recorded previously in brackish water on St. Kitts [10] and *Ae. aegypti* is tolerant of brackish water in other coastal regions [38–40]. Additionally, *Cx. quinquefasciatus* is an opportunistic forager that has the ability to fly relatively long distances [41] and so adults could be found in areas far removed from their larval breeding sites. Finally, *Ae. aegypti* was also predicted to be abundant, albeit at lower levels, in rainforest habitat across the trapping period. Elsewhere in the Caribbean, *Ae. aegypti* have been found breeding in more natural habitats in addition to artificial containers [22]. These include rock holes, calabashes, tree holes, leaf axils, bamboo joints, papaya stumps, coconut shells, bromeliads, ground pools, coral rock holes, crab holes, and conch shells which are all also present on St. Kitts.

Species with the highest capture rates and predicted by our model to have high relative abundance in mangrove habitats included both *Ae. taeniorhynchus* and *De. magnus*. These two species were predicted to have relatively low abundance in scrub surrounding mangrove sites on the island, and were not predicted to be abundant in other land cover types. *Aedes taeniorhynchus*, the black salt marsh mosquito, is widely distributed in all islands of the Caribbean where it also favors low lying marsh land as is the case on St. Kitts [21, 35]. Similarly, *De. magnus*, the crabhole mosquito, is found primarily in crabholes that are abundant in the soft sands in mangrove habitats around the Caribbean [10, 42]. We also captured *Culex nigripalpus, An. albimanus. Ps. pygmaea*, and *Aedes tortilis* in mangroves or the surrounding scrub land cover, likely due to their preference for breeding in temporary brackish and / or fresh water sources [10, 15]. We did not include these species in our relative abundance model due to low capture rates.

Generally, we had very low capture rates of mosquitoes in rainforest land cover which was consistent with our model’s predictions of a negative effect of rainforest cover on the relative abundance of the four most abundant species we trapped. This is not necessarily reflective of the results of surveys from other regions of the Caribbean. For example, a study in forested areas of eastern Trinidad between July 2007 and March 2009 collected 185,397 mosquitoes across 46 species [43]. Although this study was of a similar duration, our low capture rates in rainforest land cover might reflect 1) less breeding habitat or fewer vertebrate hosts present in the rainforest, 2) different sampling methods across the studies (e.g. CDC light traps deployed with CO_2_ lures used vs. BGS traps baited with the human lure or CDC light traps baited with sugar-yeast CO_2_ lures [12]), and 3) frequency of trapping effort (weekly vs. monthly). It seems unlikely that our trapping at ground level may have excluded mosquito species that thrive in tree-top habitats because several arboviral surveillance studies in forests in Brazil [44], New Mexico [45], and other sites across the United States [46] demonstrate that various trapping methods (e.g. entomological nets, aspirators, and CDC light traps) set in the canopy did not catch significantly more mosquitoes than those on the ground. We did capture one *Aedes busckii* in the rainforest during our survey. A previous survey [10] as well as some experience with larval surveys of tree holes in rainforest habitats on St. Kitts (data not shown) suggest *Ae. busckii* could be a rain forest habitat specialist, but more data are needed. In general, little is known about the ecology of this mosquito other than that it is confined to the Lesser Antilles (Dominica, Grenada, Guadeloupe, Martinique, Montserrat, St. Kitts and Nevis and Saint Lucia) [47]. We also captured *Cx. quinquefasciatus, Ae. taeniorhynchus*, and *De. magnus* at very low rates in the rainforest (Additional file 3: Table S3). We speculate that these captures could be the result of mosquitoes either breeding in man-made containers in neighboring agricultural habitat (*Cx. quinquefasciatus*) or mosquitoes being blown into novel habitats during tropical storms (*Ae. taeniorhynchus* and *De. magnus*). In the case of *De. magnus*, land crabs *(Gecarcinus ruricola*) do inhabit the rainforest, which could provide breeding and resting sites in their crabhole burrows for this specialist mosquito species [42].

Mosquito capture rates were also strongly determined by time of season and precipitation throughout the survey period. In general, our species count model predicted a positive effect of precipitation on the relative abundance of our four most common mosquito species. The effects of precipitation we found could be due to several reasons. Although excess rain may flush larvae from their habitats and decrease adult mosquito populations [48–49], a seasonal increase in precipitation increases the abundance and persistence of larval habitats resulting in higher densities and overall capture rates [48, 50–52]. Additionally, increased precipitation is associated with increased relative humidity, which has been shown to have important positive effects on mosquito abundance [52–53], lifespan [54], activity, and questing behavior [55–57]. Interestingly, the effect of precipitation on relative abundance was species-specific among the four most common mosquito species we found. Our post-hoc BLUPs assessment suggests the effect of precipitation had a strong effect on the relative abundances of *Ae. taeniorhynchus*, a moderate effect on *Ae. aegypti*, and smaller effects on *Cx. quinquefasciatus* and *De. magnus*. The strong effect of precipitation on *Ae. taeniorhynchus* might occur because this mosquito species relies largely on natural habitats, which are often dependent on local rainfall. While *Ae. aegypti* utilizes artificial, and human watered containers heavily for ovipositing, dependent on rainfall it is also known to oviposit in natural habitats on other Caribbean islands [22]. *Culex quinquefasciatus* might be less dependent on rainfall, as most individuals were captured in urban habitat and were most likely emerging from persistent, human-watered, artificial habitats. Whereas, increased rainfall above a certain threshold might expand water bodies in mangroves, flooding crabholes, that in turn could locally reduce breeding sites for *De. magnus* [42].

While the overall capture rates of *Ae. aegypti* were significantly lower across non-urban habitats, their presence in other land covers on St. Kitts and other Caribbean islands [10, 22] could have several implications for our understanding of the general ecology of this species and transmission of arboviruses in the Caribbean. Unlike the situation in Africa [58], where there are morphological differences between urban and non-urban forms, *Ae. aegypti* collected from urban areas of St. Kitts had the same morphology as *Ae. aegypti* collected from other land covers. Depending on the amount of gene flow that occurs across these various populations (which is thought to be limited [59–60]), ecological heterogeneity across the common land covers of St. Kitts and other Caribbean islands could result in substantial genetic variation across *Ae. aegypti* populations at fine spatial scales. Variation in microclimate [53, 61], oviposition sites [10, 22], and species of vertebrates available for blood feeding [62] are examples of potentially relevant sources of selection on *Ae. aegypti* populations inhabiting different land covers on St. Kitts. Genetic variation across *Ae. aegypti* populations on the island, in turn, could translate into substantial ecological differences among these populations with potentially significant implications for disease transmission among sylvatic reservoir hosts (e.g. non-human primates) and human populations on the island. Population genotyping is underway in our laboratories to identify if significant genetic structuring exists across *Ae. aegypti* populations on St. Kitts. Also, to more precisely determine phenotypic differences between populations, we are conducting studies to confirm the presence of reproducing adults across each land cover and using blood meal analysis and bait trapping to identify novel mosquito-host associations.

Our study provides a more comprehensive spatial and temporal (within-year) picture of the distribution of mosquito species on St. Kitts relative to previous surveys [10, 11, 15]. However, it suffers from several minor limitations. Due to specimen damage during capture and transport to the laboratory, a proportion of *Aedes* spp. (N=219) and *Culex* spp. (N=1694) were reported only to genus level. These counts differed substantially from those identified to species level which comprised 4334 *Aedes* spp. and 1697 *Culex* spp. in total. Based on the capture location of these specimens, the majority of these individuals are likely *Ae. taeniorhynchus* (mangrove) and *Cx. quinquefasciatus* (urban), respectively. That being said, capture rates for *Ae. tortilis* and *Cx. nigripalpus* are likely underestimated due to difficulty in identifying specimens damaged from capture and handling. Finally, the CDC traps performed unexpectedly poorly compared with BGS traps, possibly due to the lack of light source and the use of a human lure in the BGS traps. As trap type can select for different mosquito species [63], our count data may be biased toward species that prefer BGS traps and human lures.

## Conclusion

Consistent with findings on other islands and the literature, our island wide mosquito survey has demonstrated that the distribution and relative abundance of mosquito species on St. Kitts are affected by land cover. Further, we found substantial seasonality in mosquito capture rates, likely driven by variation in precipitation. Interesting insights gained from this study include the presence of *Ae. aegypti* in all the land covers we studied, which could have important implications for mosquito-borne disease transmission on the island. This also somewhat counters the current literature suggesting *Ae. aegypti* is primarily found in highly urban habitats and feeds almost exclusively on human hosts. Although *Aedes albopictus* occurs on other Caribbean islands [64] we did not find this species in our survey. Ongoing surveillance will be important, however, as invasive mosquito species and changing patterns of mosquito communities caused by changes in land use and climate may facilitate mosquito-borne disease transmission in people and animals.

## Additional files

Supplementary data submitted as separate files:

**Additional file 1: Table S1**. Variables and associated hypotheses evaluated in statistical models.

**Additional file 2: Table S2**. Statistical and model methods

**Additional file 3: Table S3**. Counts of mosquito species caught across the five different land covers from November 2017 to March 2019 on St. Kitts.

**Additional file 4: Table S4**. Counts of mosquito species per month on St. Kitts from Nov 2017 to March 2019 with the wet season highlighted in grey (May-November).

**Additional file 5: Figure S1**. Time series plot for each site land use category (column) and for each species (row) in their predicted relative abundance (conditional on random effects) from our best model. Solid lines denote the average predicted relative abundance of mosquito species across the 6 sites within that land cover and dotted lines denote the 95% confidence interval of that 6-site mean. Points denote the 6-site average relative abundance from the raw data. Note that the y-axis scale varies by species (row).

## Abbreviations

BGS: Biogents Sentinel 2 Trap
CO_2_: carbon dioxide
CDC: Centre for Disease Control (USA)
RUSVM: Ross University School of Veterinary Medicine
USNM: United States National Museum

## Acknowledgements

We would like to thank Gilbert Gordon, Ross University School of Veterinary Medicine Students, David Pecor, Erica McAlister, Michelle Evans, Mike Newberry, Bryan Giordano, Rob and Kathleen Gilbert, Anna Becker, Moses Humphrey, Michel Vandenplas, Jermaine Lake (Deputy Environmental Health Officer, St. Kitts and Nevis Government) and Leshan Evans, Lornette Browne, Hyacinth Richardson, Larry Greaux, Tremecia Rawlins (Vector Control Officers, St. Kitts and Nevis Government).

## Declarations

### Ethics approval and consent to participate

Not applicable

### Consent for publication

Not applicable

### Availability of data and material

The datasets supporting the conclusions of this article are included within the article and its additional files. Voucher specimens of some of the mosquito specimens associated with this study were deposited in the United States National Museum (USNM) under the following catalog numbers USNMENT01239050-74 and USNMENT01239079.

### Authors’ contributions

MV, BC and CM conducted the survey. PK and CM designed the study and supervised the survey. GJ, CA and CM produced the models. CA, MV and GJ produced tables and figures for the manuscript text. MV, CM, GJ, PK and CA prepared the manuscript text. All authors read and approved the final manuscript.

