## Additional file 1 for "Effects of seasonality and land use on the abundance and distribution of mosquitoes on St. Kitts, West Indies"

**Precip** - The sum of rainfall for the month measured at the Bradshaw International Airport.

*Expectation*: Increased rainfall is associated with higher mosquito abundance as desiccation risk of adults is reduced and viable oviposition and larval rearing sites are more abundant during periods of higher rainfall.

**m_trait** - Species-specific breeding specialization in mangrove.

*Expectation*: If a species has a mangrove breeding specialization, it is associated with higher abundances within that land cover.

**LocalAgricultural** - Percentage agricultural land cover in the 1 km area surrounding sampling sites.

*Expectation*: Nearby agricultural land cover is associated with lower abundance of non-anthrophilic mosquito species.

**LocalMangrove** - Percentage mangrove land cover in the 1 km area surrounding sampling sites.

*Expectation*: Nearby mangrove land cover is associated with higher abundance of mangrove mosquitoes.

**Mangrove** – Categorical: sampling site is located within the mangrove land cover.

*Expectation*: Mangrove land cover is associated with higher mangrove mosquito abundance within mangrove habitats.

**LocalRainforest** – Percentage rainforest land cover in the 1 km area surrounding sampling sites.

*Expectation*: Nearby rainforest land cover is associated with lower abundance of mosquitoes that are associated with mangrove or anthropogenic habitats.

**LocalUrban** - Percentage urban land cover in the 1 km area surrounding sampling sites.

*Expectation*: Nearby urban land cover is associated with higher abundances of anthrophilic mosquitoes and lower abundances of others.

**LocalScrub** – Percentage scrub land cover in the 1 km area surrounding sampling sites.

*Expectation*: Nearby scrub land cover is associated with lower mosquito abundance.

**LocalAnthropogenic** - The sum of urban and agricultural land cover.

*Expectation*: Local anthropogenic land cover is associated with higher abundances of anthrophilic mosquitoes and lower abundances of others.

**Table S1** Variables and associated hypotheses evaluated in statistical models.
