## Additional file 3 for "Effects of seasonality and land use on the abundance and distribution of mosquitoes on St. Kitts, West Indies"

**SI Table 3**: Counts of mosquito species caught across the five different land covers from November 2017 to March 2019 on St Kitts.

|  | Agricultural | Mangrove | Rainforest | Scrub | Urban | **Grand Total** |
| --- | --- | --- | --- | --- | --- | --- |
| *Aedes taeniorhynchus* | 50 | 3412 | 7 | 374 | 18 | **3861** |
| *Aedes aegypti* | 154 | 67 | 11 | 25 | 186 | **443** |
| Unidentified *Aedes* spp. | 37 | 125 | 2 | 21 | 34 | **219** |
| *Aedes tortilis* | 0 | 14 | 0 | 14 | 0 | **28** |
| *Aedes busckii* | 1 | 0 | 1 | 0 | 0 | **2** |
| *Anopheles albimanus* | 1 | 3 | 0 | 0 | 0 | **4** |
| *Culex quinquefasciatus* | 23 | 261 | 1 | 566 | 812 | **1663** |
| Unidentified *Culex* spp. | 40 | 258 | 3 | 105 | 1288 | **1694** |
| *Culex nigripalpus* | 0 | 15 | 0 | 0 | 19 | **34** |
| *Culex bahamensis* | 0 | 0 | 0 | 0 | 0 | **0** |
| *Culex bisulcatus* | 0 | 0 | 0 | 0 | 0 | **0** |
| *Culex declarator* | 0 | 0 | 0 | 0 | 0 | **0** |
| *Culex madininensis* | 0 | 0 | 0 | 0 | 0 | **0** |
| *Deinocerites magnus* | 1 | 1533 | 8 | 26 | 9 | **1577** |
| *Psorophora pygmaea* | 0 | 0 | 0 | 178 | 0 | **178** |
| *Toxorhynchites guadeloupensis* | 0 | 0 | 0 | 1 | 0 | **1** |
| **Grand Total** | **307** | **5688** | **33** | **1310** | **2366** | **9704** |
