## Additional file 4 for "Effects of seasonality and land use on the abundance and distribution of mosquitoes on St. Kitts, West Indies"

|  | Nov-17 | Jan-18 | Feb-18 | Mar-18 | May-18 | Jun-18 | Jul-18 | Aug-18 | Sep-18 | Oct-18 | Nov-18 | Jan-19 | Feb-19 | Mar-19 | Grand Total |
| --- | --- | --- | --- | --- | --- | --- | --- | --- | --- | --- | --- | --- | --- | --- | --- |
| *Aedes taeniorhynchus* | 63 | 89 | 18 | 6 | 0 | 9 | 321 | 5 | 111 | 811 | 2377 | 14 | 9 | 28 | 3861 |
| *Aedes aegypti* | 50 | 9 | 6 | 10 | 6 | 20 | 24 | 26 | 41 | 147 | 80 | 11 | 11 | 2 | 443 |
| Unidentified *Aedes* spp. | 71 | 24 | 0 | 1 | 9 | 4 | 10 | 9 | 34 | 41 | 16 | 0 | 0 | 0 | 219 |
| *Aedes tortilis* | 0 | 0 | 0 | 0 | 0 | 0 | 0 | 0 | 0 | 0 | 28 | 0 | 0 | 0 | 28 |
| *Aedes busckii* | 0 | 0 | 0 | 0 | 0 | 0 | 0 | 1 | 0 | 0 | 0 | 1 | 0 | 0 | 2 |
| *Anopheles albimanus* | 1 | 0 | 0 | 3 | 0 | 0 | 0 | 0 | 0 | 0 | 0 | 0 | 0 | 0 | 4 |
| *Culex quinquefasciatus* | 37 | 0 | 10 | 37 | 34 | 36 | 396 | 69 | 77 | 315 | 244 | 114 | 153 | 141 | 1663 |
| Unidentified *Culex* spp. | 210 | 19 | 21 | 63 | 9 | 22 | 27 | 98 | 360 | 544 | 318 | 3 | 0 | 0 | 1694 |
| *Culex nigripalpus* | 14 | 0 | 1 | 0 | 0 | 0 | 0 | 0 | 0 | 2 | 17 | 0 | 0 | 0 | 34 |
| *Culex bahamensis* | 0 | 0 | 0 | 0 | 0 | 0 | 0 | 0 | 0 | 0 | 0 | 0 | 0 | 0 | 0 |
| *Culex bisulcatus* | 0 | 0 | 0 | 0 | 0 | 0 | 0 | 0 | 0 | 0 | 0 | 0 | 0 | 0 | 0 |
| *Culex declarator* | 0 | 0 | 0 | 0 | 0 | 0 | 0 | 0 | 0 | 0 | 0 | 0 | 0 | 0 | 0 |
| *Culex madininensis* | 0 | 0 | 0 | 0 | 0 | 0 | 0 | 0 | 0 | 0 | 0 | 0 | 0 | 0 | 0 |
| *Deinocerites magnus* | 11 | 0 | 3 | 14 | 1 | 38 | 104 | 179 | 284 | 170 | 531 | 21 | 45 | 176 | 1577 |
| *Psorophora pygmaea* | 0 | 0 | 0 | 0 | 1 | 1 | 0 | 0 | 1 | 0 | 175 | 0 | 0 | 0 | 178 |
| *Toxorhynchites guadeloupensis* | 0 | 0 | 0 | 1 | 0 | 0 | 0 | 0 | 0 | 0 | 0 | 0 | 0 | 0 | 1 |
| Grand Total | 457 | 141 | 59 | 135 | 60 | 130 | 882 | 387 | 908 | 2030 | 3786 | 164 | 218 | 347 | 9704 |

**SI Table 4:** Counts of mosquito species per month on St Kitts from Nov 2017 to March 2019 with the wet season highlighted in grey (May-November).
