## Supplementary figures and images for "Effects of seasonality and land use on the abundance and distribution of mosquitoes on St. Kitts, West Indies"

### Additional file 5

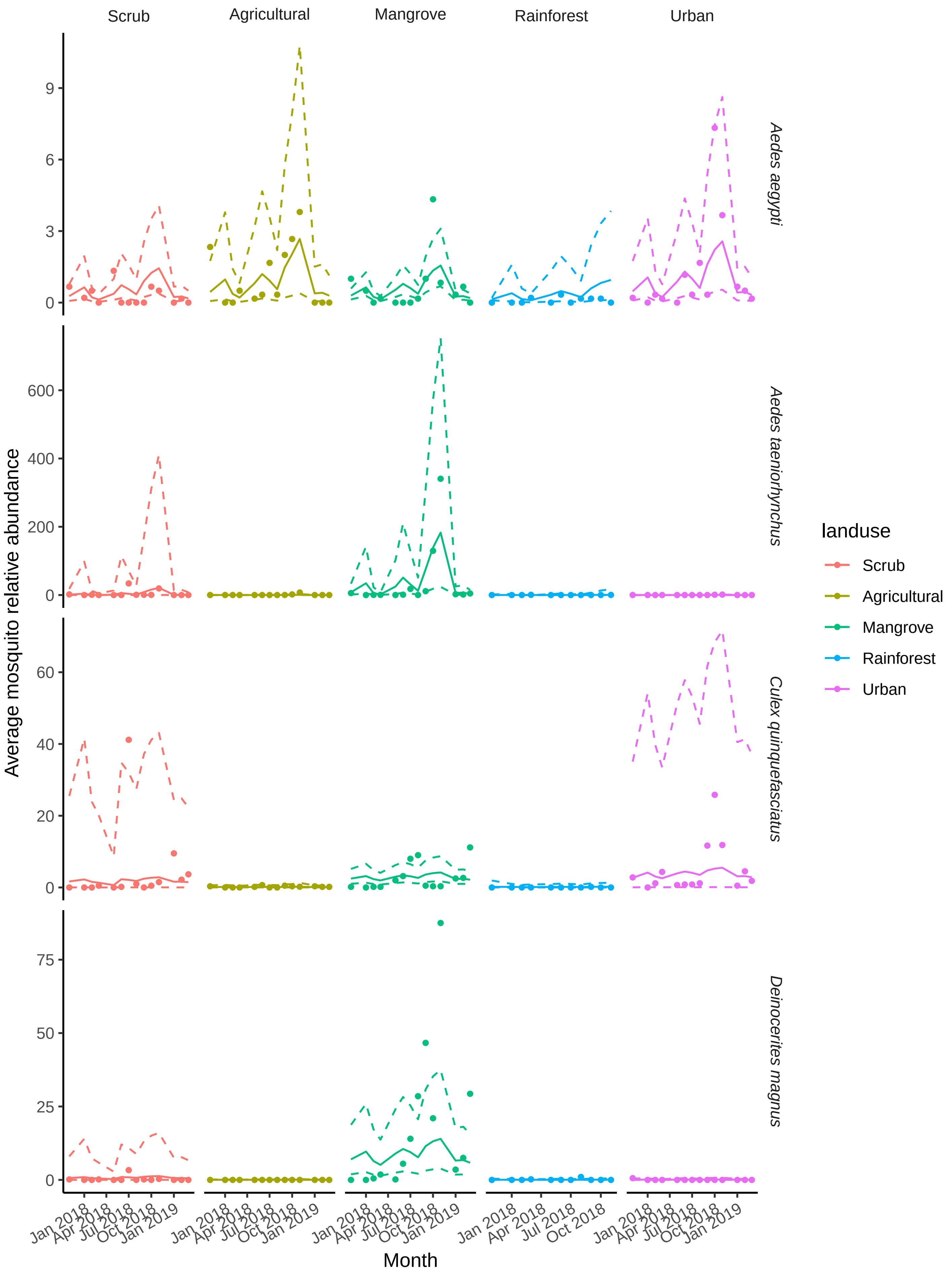
